## Supplementary information for "Rewiring capsule production by CRISPRi-based genetic oscillators demonstrates a functional role of phenotypic variation in pneumococcal-host interactions"

### Strain construction:

**Strain VL3251 (D39V, *ΔPrs1::*$P_{F6}$-*lacI* (*gen*), *bgaA*::$P_{lac}$-*dCas9sp* (*tet*), *cil*::$P_3$-*luc* (*kan*), *cep*::$P_3$-BS1-sgRNA*luc* (*spc*))** was constructed by Golden Gate assembly of two parts as follows. The first part was amplification by PCR of the upstream homologous region to integrate in the *S. pneumoniae* (*S.p.*) *cep* locus, the *spc* marker, the promoter $P_3$ and BS1', from plasmid pVL1305 with primers OVL3205 (with BsmBI restriction site) and OVL2625. Second part is the BS1', the base pairing *luc*, the terminator *S. pyogenes* and the downstream homologous region to integrate in *S. p. cep* locus, amplified by PCR from the plasmid pVL1305 with primers OVL3204 (with BsmBI restriction site) and OVL726. After purification, the two parts were digested with BsmBI (NEB) at 55°C for 4h. After purification, ligation was realized at room temperature with T4 DNA ligase (Vazyme) 1h and directly used to transform *S.p.* strain VL997 with spectinomycin selection. The *cep* locus of the resulting strain, VL3252 ,was confirmed by Sanger sequencing.

**Strain VL3252 (D39V, *ΔPrs1::*$P_{F6}$-*lacI* (*gen*), *bgaA*::$P_{lac}$-*dCas9sp* (*tet*), *cil*::$P_3$-*luc* (*kan*), *cep*::$P_3$- BS2-sgRNA*luc* (*spc*))**, was constructed by Golden Gate assembly of two parts identical to the construction of strain VL3251. First part is the amplification by PCR of pVL1305 with primers OVL3207 (with BsmBI restriction site) and OVL2625. Second part is the amplification by PCR of pVL1305 with primers OVL3206 (with BsmBI restriction site) and OVL726.

**Strain VL3253 (D39V, *Δprs1*::$P_{F6}$-*lacI* (*gen*), *bgaA*::$P_{lac}$-*dCas9sp* (tet), *cil*::$P_3$-*luc* (*kan*), *cep*::P3- BS3-sgRNA*luc* (*spc*))**, was constructed by Golden Gate assembly of two parts identical to the construction of strain VL3251. First part is the amplification by PCR of pVL1305 with primers OVL3385 (with BsmBI restriction site) and OVL2625. Second part is the amplification by PCR of pVL1305 with primers OVL3384 (with BsmBI restriction site) and OVL726.

**Strain VL3254 (D39V, *ΔPrs1::*$P_{F6}$-*lacI* (*gen*), *bgaA*::$P_{lac}$-*dCas9sp* (*tet*), *cil*::$P_3$-*luc* (*kan*), *cep*::$P_3$- BS4-sgRNA*luc* (*spc*))**, was constructed by Golden Gate assembly of two parts identical to the construction of strain VL3251. First part is the amplification by PCR of pVL1305 with primers OVL3209 (with BsmBI restriction site) and OVL2625. Second part is the amplification by PCR of pVL1305 with primers OVL3208 (with BsmBI restriction site) and OVL726.

**Strain VL3255 (D39V, *ΔPrs1::*$P_{F6}$-*lacI* (*gen*), *bgaA*::$P_{lac}$-*dCas9sp* (*tet*), *cil*::$P_3$-*luc* (*kan*), *cep*::$P_3$- BS5-sgRNA*luc* (*spc*))**, was constructed by Golden Gate assembly of two parts identical to the construction of strain VL3251. First part is the amplification by PCR of pVL1305 with primers OVL3387 (with BsmBI restriction site) and OVL2625. Second part is the amplification by PCR of pVL1305 with primers OVL3386 (with BsmBI restriction site) and OVL726.

**Strain VL3256 (D39V, *Δprs1*::$P_{F6}$-*lacI* (gm), *bgaA*::$P_{lac}$-*dCas9sp* (tet), *cil*::$P_3$-*luc* (kan), *cep*::$P_3$- BS6-sgRNA*luc* (spc))** was constructed by Golden Gate assembly of two parts identical to the construction of strain VL3251. First part is the amplification by PCR of pVL1305 with primers OVL3389 (with BsmBI restriction site) and OVL2625. Second part is the amplification by PCR of pVL1305 with primers OVL3388 (with BsmBI restriction site) and OVL726.

**Strain VL3308 (D39V, *Δprs1*::$P_{F6}$-*lacI/tetR* (*gen*), *bgaA*::$P_{lac}$-*dCas9sp* (*tet*))** was constructed by amplification by PCR of the upstream homologous region to integrate in the *S. p. bgaA* locus, the *tet* marker, $P_{lac}$-*dCas9sp* and the downstream homologous region using chromosomal DNA (gDNA) of strain VL1998 as template with primers OVL173 and OVL174. After purification, the PCR product was used to transform strain VL333 with tetracycline selection. The *bgaA* locus of the resulting strain, VL3308, was confirmed by Sanger sequencing.

**Strain VL3436 (D39V, *Δprs1*::$P_{F6}$-*lacI/tetR* (*gen*), *bgaA*::$P_{lac}$-*dCas9sp* (*tet*), *cep*::$P_3$-BS3-sgRNA2 (*spc*))** was constructed by Golden Gate assembly of two parts. First part is the amplification by PCR of the upstream homologous region to integrate in the *S. p. cep* locus, the *spc* marker, the promoter $P_3$ and the BS3 from the gDNA of strain VL3253 with primers OVL3504 (with BsmBI restriction site) and OVL2625. Second part is the amplification by PCR of the sgRNA2, the *dCas9* handle, the sgRNA terminator and the downstream homologous region to integrate in *S. p. cep* locus, from the gDNA of strain VL3253 with OVL3205 (with BsmBI restriction site) and OVL2628. After purification, the two parts were digested with BsmBI (NEB) at 55°C for 2h. After purification, ligation was realized at room temperature with T4 DNA ligase (Vazyme) for 1h and directly used to transform strain VL3308 with spectinomycin selection. The *cep* locus of the resulting strain, VL3436, was confirmed by Sanger sequencing.

**Strain VL3437 (D39V, *Δprs1*::$P_{F6}$-*lacI/tetR* (*gen*), *bgaA*::$P_{lac}$-*dCas9sp* (*tet*), *cep*::$P_3$-BS6-sgRNA3 (*spc*))** was constructed by Golden Gate assembly of two parts. First part is the amplification by PCR of the upstream homologous region to integrate in *S. p. cep* locus, the *spc* marker, the promoter $P_3$ and the BS3 from the gDNA of strain VL3256 with the primers OVL3506 (with BsmBI restriction site) and OVL2625. Second part is the amplification by PCR of the sgRNA3, the dCas9 handle, the sgRNA terminator and the downstream homologous region to integrate in *S. p. cep* locus, from the gDNA of strain VL3256 with primers OVL3207 (with BsmBI restriction site) and OVL2628. After purification, the two parts were digested with BsmBI (NEB) at 55°C for 2h. After purification, ligation was realized at room temperature with T4 DNA ligase (Vazyme) for 1h and directly used to transform strain VL3308 with spectinomycin selection. The *cep* locus of the resulting strain, VL3437 was confirmed by Sanger sequencing.

**Strain VL3438 (D39V, *Δprs1*::$P_{F6}$-*lacI/tetR* (*gen*), *bgaA*::$P_{lac}$-*dCas9sp* (*tet*), *cep*::$P_3$-BS2-sgRNA6 (*spc*))** was constructed by Golden Gate assembly of two parts. First part is the amplification by PCR of the upstream homologous region to integrate in *S. p. cep* locus, the *spc* marker, the promoter $P_3$ and the BS2 from the gDNA of strain VL3252 with the primers OVL3502 (with BsmBI restriction site) and OVL2625. Second part is the amplification by PCR of the sgRNA6, the dCas9 handle, the sgRNA terminator and the downstream homologous region to integrate in the *S. p. cep* locus, from the gDNA of strain VL3252 with OVL3203 (with BsmBI restriction site) and OVL2628. After purification, the two parts were digested with BsmBI (NEB) at 55°C for 2h. After purification, ligation was realized at room temperature with T4 DNA ligase (Vazyme) for 1h and directly used to transform strain VL3308 with spectinomycin selection. The *cep* locus of the resulting strain, VL3438 was confirmed by Sanger sequencing.

**Strain VL3439 (D39V, *Δprs1*::P$_{F6}$-*lacI/tetR* (*gen*), *bgaA*::P$_{lac}$-*dCas9sp* (*tet*), *cep*::P$_3$-BS3-sgRNA2 (*spc*), *zip*::P$_3$-BS2-*mNeonGreen-opt* (*ery*))** was constructed by Golden Gate assembly of three parts. First part is the amplification by PCR of the upstream homologous region to integrate in the *S. p. zip* locus and the *ery* marker from the plasmid pASR103 with primers OVL3531 and OVL3851 (with BsmBI restriction site). Second part is the amplification by PCR of T_rpsI, T_tuf, T_B1002, T_B0015, the promoter P$_3$ and the BS2 from the gDNA of strain VL3252 with primers OVL3533 and OVL3534 (both with BsmBI restriction site). Third part is the amplification by PCR of *mNeonGreen- optimized* and the downstream homologous region to integrate in *S. p. zip* locus from the plasmid pASR110 with primers OVL3535 (with BsmBI restriction site) and OVL3536. After DpnI treatment for parts 1 and 3 and purification, the three parts were digested with BsmBI (NEB) at 55°C for 3h. After purification, ligation was realized at room temperature with T4 DNA ligase (Vazyme) for 1h and directly used to transform strain VL3436 with erythromycin selection. The *zip* locus of the resulting strain, VL3439 was confirmed by Sanger sequencing.

**Strain VL3440 (D39V, *Δprs1*::P$_{F6}$-*lacI/tetR* (*gen*), *bgaA*::P$_{lac}$-*dCas9sp* (*tet*), *cep*::P$_3$-BS3-sgRNA2 (*spc*), *zip*::P$_3$-BS2-*mNeonGreen-opt* (*ery*))** was constructed by Golden Gate assembly of three parts. First part is the amplification by PCR of the upstream homologous region to integrate in *S. p. zip* locus and the *ery* marker from plasmid pASR103 with primers OVL3531 and OVL3851 (with BsmBI restriction site). Second part is the amplification by PCR of T_rpsI, T_tuf, T_B1002, T_B0015, the promoter P$_3$ and the BS6 from gDNA of strain VL3253 with primers OVL3533 and OVL3537 (both with BsmBI restriction site). Third part is the amplification by PCR of *mTurquoise2-optimized* and the downstream homologous region to integrate in *S. p. zip* locus from gDNA of template strain VL3312 (Veening lab collection) with primers OVL3538 (with BsmBI restriction site) and OVL3536. After DpnI treatment of part 1 and purification, the three parts were digested with BsmBI (NEB) at 55C for 3h. After purification, ligation was realized at room temperature with T4 DNA ligase (Vazyme) for 1h and directly used to transform strain VL3437 with erythromycin selection. The *zip* locus of the resulting strain, VL3440, was confirmed by Sanger sequencing.

**Strain VL3441 (D39V, *Δprs1*::P$_{F6}$-*lacI/tetR* (*gen*), *bgaA*::P$_{lac}$-*dCas9sp* (*tet*), *cep*::P$_3$-BS2-sgRNA6 (*spc*), *zip*::P$_3$-BS6-*mScarletI-opt* (*ery*))** was constructed by Golden Gate assembly of three parts. First part is the amplification by PCR of the upstream homologous region to integrate in *S. p. zip* locus and the *ery* marker from plasmid pASR103 with primers OVL3531 and OVL3851 (with BsmBI restriction site). Second part is the amplification by PCR of T_rpsI, T_tuf, T_B1002, T_B0015, the promoter P$_3$ and the BS2 from gDNA of strain VL3256 with primers OVL3533 and OVL3539 (both with BsmBI restriction site). Third part is the amplification by PCR of *mScarletI-optimized* and the downstream homologous region to integrate in *S. p. zip* locus from the template strain VL3309 (Veening lab collection) with primers OVL3540 (with BsmBI restriction site) and OVL3536. After DpnI treatment of part 1 and purification, the three parts were digested with BsmBI (NEB) at 55°C for 3h. After purification, ligation was realized at room temperature with T4 DNA ligase (Vazyme) for 1h and directly used to transform strain VL3438 with erythromycin selection. The *zip* locus of the resulting strain, VL3441, was confirmed by Sanger sequencing.

**Strain VL3746 (D39V, *Δprs1*::P$_{F6}$-*lacI/tetR* (*gen*), *bgaA*::P$_{lac}$-*dCas9sp* (*tet*), *zip*::P$_3$-BS6-sgRNA3-P$_3$-BS2-*mNeonGreen-opt* (*ery*))** was constructed by Golden Gate assembly of three parts. First part is the amplification by PCR of the upstream homologous region to integrate in *S. p. zip* locus, the *ery* marker and the terminators T_rpsI, T_tuf from the template strain VL3441 with primers OVL3252 and OVL4250 (with BsmBI restriction site). Second part is the amplification by PCR of P$_3$, BS6, sgRNA3, the dCas9 handle and the sgRNA terminator using gDNA of strain VL3437 as template with OVL4251 and OVL4252 (both with BsmBI restriction site). Third part is the amplification by PCR of terminators T_B1002, T_B0015, P$_3$, BS2, *mNeonGreen-optimized*, T_B1006 and the downstream homologous region to integrate in *S. p. zip* locus from the template strain VL3439 with OVL4253 (with BsmBI restriction site) and OVL3255. After purification, the three parts were digested and ligated together during an assembly reaction with T4 DNA ligase buffer (Vazyme), T4 DNA ligase (Vazyme) and Eps3i (NEB; isoschizomer of BsmBI). The assembly mixture was incubated in a thermocycler (PCR Max) for step 1: 1.5 min at 37°C, step 2: 3 min at 16°C, steps 1 and 2 repeated 25x, step 3: 5 min at 37°C and step 4: 10 min at 80°C. Transformation of VL3308 with the assembly mixture and selection with erythromycin. The *zip* locus of the resulting strain, VL3746, was confirmed by Sanger sequencing.

**Strain VL3747 (D39V, *Δprs1*::$P_{F6}$-*lacI/tetR* (*gen*), *bgaA*::$P_{lac}$-*dCas9sp* (*tet*), *cep*::$P_3$-BS6-*mScarletI-opt*-$P_3$-BS3-sgRNA2 (*spc*))** was constructed by Golden Gate assembly of three parts. First part is the amplification by PCR of the upstream homologous region to integrate in *S. p. cep* locus, the *spc* marker and the terminators T_rpsI, T_tuf from the template strain VL3436 with the primers OVL725 and OVL4254 (with BsmBI restriction site). Second part is the amplification by PCR of $P_3$, BS6 and *mScarletI-optimized* from the template strain VL3441 with primers OVL4255 and OVL4256 (both with BsmBI restriction site). Third part is the amplification by PCR of terminators T_B1002, T_B0015, $P_3$, BS3, sgRNA2, the *dCas9 handle*, the terminator *S. pyogenes*, T_B1006 and the downstream homologous region to integrate in *S. p. cep* locus from the template strain VL3436 with primers OVL4257 (with BsmBI restriction site) and OVL726. After purification, the three parts were digested and ligated together during an assembly reaction with T4 DNA ligase buffer (Vazyme), T4 DNA ligase (Vazyme) and Eps3i (NEB). The assembly mixture was incubated in a thermocycler (PCR Max) for step 1: 1.5 min at 37°C, step 2: 3 min at 16°C, steps 1 and 2 repeated 25x, step 3: 5 min at 37°C and step 4: 10 min at 80°C. Transformation of VL3308 with the assembly mixture and selection with erythromycin. The *cep* locus of the resulting strain, VL3747, was confirmed by Sanger sequencing.

**Strain VL3748 (D39V, *Δprs1*::$P_{F6}$-*lacI/tetR* (*gen*), *bgaA*::$P_{lac}$-*dCas9sp* (*tet*), *cil*::$P_3$-BS3-*mTurquoise2-opt*-$P_{tet}$-BS2-sgRNA6 (*kan*))** was constructed by Golden Gate assembly of five parts. First part is the amplification by PCR of the upstream homologous region to integrate in *S. p. cil* locus, the *kan* marker and MCS from the template strain VL2969 with the primers OVL3318 and OVL4258 (with BsmBI restriction site). Second part is the amplification by PCR of $P_3$, BS3 and *mTurquoise2-opt* from the template strain VL3440 with primers OVL4259 and OVL4260 (both with BsmBI restriction site). Third part is the amplification by PCR of terminators T_rrnB and T_rpsI from the template strain VL2969 with primers OVL4261 and OVL4262 (both with BsmBI restriction site). The fourth part is the amplification by PCR of BssgRNA2, sgRNA-6 the *dCas9 handle*, the sgRNA terminator from template strain VL3438 with primers OVL4263 and OVL4264 (both with BsmBI restriction site). Fifth part is the amplification by PCR of the terminator T_tufA and the downstream homologous region to integrate in *S. p. cil* locus from the template strain VL2969 with the primers OVL4265 (with BsmBI restriction site) and OVL4266. After purification, the five parts were digested and ligated together during an assembly reaction with T4 DNA ligase buffer (Vazyme), T4 DNA ligase (Vazyme) and Eps3i. The assembly mixture was incubated in a thermocycler (PCR Max) for step 1: 1.5 min at 37°C, step 2: 3 min at 16°C, steps 1 and 2 repeated 25x, step 3: 5 min at 37°C and step 4: 10 min at 80°C. The assembly mixture was amplified by PCR with OVL3318 and OVL4266. After purification by gel extraction, the product was used to transform strain VL3308 and selection with kanamycin. The *cil* locus of the resulting strain, VL3748, was confirmed by Sanger sequencing. This construct has a point mutation in the *mTurquoise2-opt* gene (leading to a F72I amino acid change) as well as an insertion of a second MCS sequence between the T_tufA and the downstream homologous region to integrate in *S. p. cil* locus.

**Strain VL3749 (D39V, *Δprs1*::$P_{F6}$-*lacI/tetR* (*gen*), *bgaA*::$P_{lac}$-*dCas9sp* (*tet*), *cil*::$P_3$-BS3-*mTurquoise2-opt*-$P_3$-BS2-sgRNA6 (*kan*))** was constructed by Golden Gate assembly of five parts in the same fashion as the strain VL3748. The first, second and fifth parts are similar as the one of VL3748. Parts three and four are different as described. Third part was amplification by PCR of terminators T_rrnB and T_rpsI from the template strain VL2969 with primers OVL4261 and OVL4267 (both with BsmBI restriction site). The fourth part was amplification by PCR of BS2, sgRNA6, *dCas9 handle* and sgRNA Terminator from template strain VL3438 with primers OVL4268 and OVL4264 (both with BsmBI restriction site). This construct has an insertion of a second MCS sequence between the T_tufA and the downstream homologous region to integrate in *S. p. cil* locus.

**Strain VL3750 (D39V, *Δprs1::*$P_{F6}$*-lacI/tetR* (*gen*), *bgaA*::$P_{lac}$-*dCas9sp* (*tet*), *zip*::$P_3$-BS6-sgRNA3-$P_3$-BS2-*mNeonGreen-opt* (*ery*), *cep*::$P_3$-BS6-*mScarletI-opt*-$P_3$-BS3-sgRNA2 (*spc*))** was constructed by amplification by PCR of the *cep* locus of the strain VL3747 with the primers OVL3340 and OVL3341 (both without restriction site). After purification, VL3746 was transformed with spc selection. The *cep* locus of the resulting strain, VL3750, was confirmed by Sanger sequencing.

**Strain VL3752 (D39V, *Δprs1::*$P_{F6}$*-lacI/tetR* (*gen*), *bgaA*::$P_{lac}$-*dCas9sp* (*tet*), *zip*::$P_3$-BS6-sgRNA3-$P_3$-BS2-*mNeonGreen-opt* (*ery*), *cep*::$P_3$-BS6-*mScarletI-opt*-$P_3$-BS3-sgRNA2 (*spc*), *cil*::$P_3$-BS3-*mTurquoise2-opt*-$P_{tet}$-BS2-sgRNA6 (*kan*))** was constructed by amplification by PCR of the *cil* locus of strain VL3749 with primers OVL552 and OVL553 (both without restriction site). After purification by gel extraction, VL3750 was transformed with this product while selecting on agar plates containing kanamycin. The *cil* locus of the resulting strain, VL3752, was confirmed by Sanger sequencing.

**Strain VL3753 (D39V, *Δprs1::*$P_{F6}$*-lacI/tetR* (*gen*), *bgaA*::$P_{lac}$-*dCas9sp* (*tet*), *zip*::$P_3$-BS6-sgRNA3-$P_3$-BS2-*mNeonGreen-opt* (*ery*), *cep*::$P_3$-BS6-*mScarletI-opt*-$P_3$-BS3-sgRNA2 (*spc*), *cil*::$P_3$-BS3-*mTurquoise2-opt*-$P_3$-BS2-sgRNA6 (*kan*))** was constructed by amplification by PCR of the *cil* locus of strain VL3749 with primers OVL1289 and OVL3321 (both without restriction site). After purification by gel extraction, VL3750 was transformed with kan selection. The *cil* locus of the resulting strain, VL3753, was confirmed by sequencing.

**Strain VL3755 (D39V, *Δprs1*:: $P_{F6}$*-lacI/tetR* (*gen*), *bgaA*::$P_{lac}$-*dCas9*sp (*tet*), *zip*::$P_3$-BS6-sgRNA3-$P_3$-BS2-*mNeonGreen-opt* (*ery*), *cep*::$P_3$-BS6-*mScarletI-opt*-$P_3$-BS3-sgRNA2 (*spc*), *cil*::$P_3$-BS3-*mTurquoise2-opt*-$P_{tet}$-BS2-sgRNA6 (*kan*), *lytA::cat*, *comC::tmp*)** is strain VL3752 with deletion of *lytA* and *comC* genes. Deletion of *lytA* was realized by amplification of the *lytA::cat* region from gDNA of strain *lytA::cat* (lab collection) with OVL3171 and OVL3172 and purified. In parallel, the deletion of *comC::tmp* was realized by Golden Gate assembly of 3 parts. The flanking regions were amplified by PCR from strain VL783. Primers OVL4661 and OVL4662 (with BsmBI restriction site) were used for the upstream insertion site. Primers OVL4665 and OVL4666 (with BsmBI restriction site) were used for the downstream insertion site. The *tmp* marker was amplified by PCR from gDNA of strain VL3882 with OVL4663 and OVL4664 (both with BsmBI restriction site). After purification, the three parts were digested and ligated together during an assembly reaction with T4 DNA ligase buffer (Vazyme), T4 DNA ligase (Vazyme) and Eps3i (NEB). The assembly mixture was incubated in a thermocycler (BioRad) for step 1: 1.5 min at 37°C, step 2: 3 min at 16°C, steps 1 and 2 repeated 25x, step 3: 5 min at 37°C and step 4: 10 min at 80°C. The assembly mixture was purified by gel extraction. VL3752 was transformed by both the PCR product of *lytA::cat* and the purified assembly product of *comC::tmp*, with double selection, chloramphenicol (cm) and tmp. The *lytA* and *comC* loci of the resulting strain, VL3755, were confirmed by Sanger sequencing.

**Strain VL3757 (D39V, *Δprs1*::$P_{F6}$*-lacI/tetR* (*gen*), *bgaA*::$P_{lac}$-*dCas9sp* (*tet*), *zip*::$P_3$- BS6-sgRNA3-$P_3$-BS2-*mNeonGreen-opt* (*ery*), *cep*::$P_3$- BS6-*mScarletI-opt*-$P_3$-BS3-sgRNA2 (*spc*), *cil*::$P_3$-BS3-*mTurquoise2-opt*-$P_3$-BS2-sgRNA6 (*kan*), *lytA::cat*, *comC::tmp*)** is VL3753 with deletion of *lytA* and *comC* genes. The double deletion was realized as described for VL3755.

**Strain VL3869 (D39V, *Δprs1*:: $P_{F6}$-*lacI/tetR* (*gen*), *bgaA*::Plac-*dCas9*sp (*tet*), *zip*::$P_3$-BS6-sgRNA3-$P_3$-BS2-*mNeonGreen-opt* (*ery*), *cep*::$P_3$-BS6-*mScarletI-opt*-$P_3$-BS3-sgRNA2 (*spc*), *cil*::$P_3$-BS3-*mTurquoise2-opt*-$P_{tet}$-BS2-sgRNA6 (*kan*), *lytA::cat*, *comC::tmp, Δcps*)** is VL3755 with deletion of the *cps* operon without an antibiotic marker. The region *Δcps* was amplified by PCR from strain VL3660 with primers OVL4933 and OVL4938. After purification, VL3755 was transformed without antibiotic selection and 10 colonies were screened for absence of capsule. One of the positive clones lacking capsule was further analyzed and confirmed by Sanger sequencing resulting in strain VL3869.

**Strain VL3870 (D39V, *bgaA::*$P_{lac}$-*dcas9sp* (*gen*))** was constructed by Golden Gate assembly to replace the *tet* marker from VL3753 to *gen* marker. The flanking regions were amplified by PCR from strain VL3753 with primers OVL5139 and OVL5140 (with SapI restriction site) for the upstream and OVL5143 (with SapI restriction site) and OVL5144 for the downstream. The *gen* marker was amplified from pASR102 with OVL5141 and OVL5142 (both with SapI restriction site). After purification, the three parts were digested and ligated together during an assembly reaction with T4 DNA ligase buffer (Vazyme), T4 DNA ligase (Vazyme) and SapI. The assembly mixture was incubated in a thermocycler (PCR Max) for step 1: 1.5 min at 37°C, step 2: 3 min at 16°C, steps 1 and 2 repeated 25x, step 3: 5 min at 37°C and step 4: 10 min at 80°C. Strain VL1 was directly transformed with the assembly product with gentamycin selection. The *bgaA* locus of the resulting strain, VL3870, was confirmed by Sanger sequencing.

**Strain VL3872 (D39V, $P_{cps}$::$P_3$-BS6-*mScarletI-opt-cps* (*tet*))** was constructed by Golden Gate assembly. The flanking regions were amplified by PCR from the strain VL3703 with the primers OVL5145 and OVL5146 (with SapI restriction site) for the upstream and OVL5149 (with SapI restriction site) and OVL5150 for the downstream. $P_3$-BS6-*mScarletI-opt* was amplified from strain VL3747 with primers OVL5147 and OVL5148 (both with SapI restriction site). The assembly mixture was incubated in a thermocycler (PCR Max) as described for VL3871. Strain VL1 was directly transformed with the assembly product with tetracycline selection. The *cps* locus of the resulting strain, VL3872, was confirmed by Sanger sequencing.

**Strain VL3873 (D39V, $P_{cps}$::$P_3$-BS6-*mScarletI-opt-cps* (*tet*), *cep*::$P_3$-BS3-sgRNA2 (*spc*))** was constructed by amplification of *cep* locus of strain VL3436 with the primers OVL725 and OVL726, the PCR product purified was used to transform VL3872 with spc selection. The *cep* locus of the resulting strain, VL3873, was confirmed by Sanger sequencing.

**Strain VL3875 (D39V, $P_{cps}$::$P_3$-BS6-*mScarletI-opt-cps* (*tet*), *cep*::$P_3$-BS3-sgRNA2 (*spc*), *zip*::$P_3$-BS6-sgRNA3-$P_3$-BS2-*mNeonGreen-opt* (*ery*))** was constructed by amplification of the *zip* locus of strain VL3746 with primers OVL3531 and OVL3536, the PCR product purified was used to transform VL3873 with ery selection. The *zip* locus of the resulting strain, VL3875, was confirmed by colony PCR.

**Strain VL3876 (D39V, $P_{cps}$::$P_3$-BS6-*mScarletI-opt-cps* (*tet*), *cep*::$P_3$-BS3-sgRNA2 (*spc*), *zip*::$P_3$-BS6-sgRNA3-$P_3$-BS2-*mNeonGreen-opt* (*ery*), *cil*::$P_3$-BS3-*mTurquoise2-opt*-$P_{tet}$-BS2-sgRNA6 (*kan*))** was constructed by amplification of the *cil* locus of strain VL3752 with primers OVL3318 and OVL3321, the PCR product purified was used to transform VL3875 with kan selection. The *cil* locus of the resulting strain, VL3876, was confirmed by Sanger sequencing.

**Strain VL3877 (D39V, $P_{cps}$::$P_3$-BS6-*mScarletI-opt-cps* (*tet*), *cep*::$P_3$-BS3-sgRNA2 (*spc*), *zip*::$P_3$-BS6-sgRNA3-$P_3$-BS2-*mNeonGreen-opt* (*ery*), *cil*::$P_3$-BS3-*mTurquoise2-opt*-$P_3$-BS2-sgRNA6 (*kan*))** was constructed by amplification of the *cil* locus of the strain VL3753 with primers OVL3318 and OVL3321, the purified PCR product was used to transform VL3875 with kan selection. The *cil* locus of the resulting strain, VL3877, was confirmed by Sanger sequencing.

**Strain VL3878 (D39V, $P_{cps}$::$P_3$-BS6-*mScarletI-opt-cps* (*tet*), *cep*::$P_3$-BS3-sgRNA2 (*spc*), *zip*::$P_3$-BS6-sgRNA3-$P_3$-BS2-*mNeonGreen-opt* (*ery*), *cil*::$P_3$-BS3-*mTurquoise2-opt*-$P_{tet}$-BS2-sgRNA6 (*kan*), *bgaA*::$P_{lac}$-*dcas9sp* (*gen*))** was constructed by amplification of the *bgaA* locus of strain VL3870 with primers OVL173 and OVL174, the PCR product purified was used to transform VL3876 with gentamycin selection. The resulting strain, VL3878, was confirmed by colony PCR and microscopy.

**Strain VL3879 (D39V, $P_{cps}$::$P_3$-BS6-*mScarletI-opt-cps* (*tet*), *cep*::$P_3$-BS3-sgRNA2 (*spc*), *zip*::$P_3$-BS6-sgRNA3-$P_3$-BS2-*mNeonGreen-opt* (*ery*), *cil*::$P_3$-BS3-*mTurquoise2-opt*-$P_3$-BS2-sgRNA6 (*kan*), *bgaA*::$P_{lac}$-*dcas9sp* (*gen*))** was constructed by amplification of the *bgaA* locus of strain VL3870 with primers OVL173 and OVL174. The purified PCR product was used to transform VL3877 with gentamycin selection. The resulting strain, VL3879, was confirmed by colony PCR and microscopy.

**Strain VL4313 (D39V, *zip*::$P_3$- BS6-sgRNA3- $P_3$-BS2-*mNeonGreen-opt* (*ery*))** was constructed by amplification of the *zip* locus of strain VL3746 with primers OVL3531 and OVL3536. The purified PCR product was used to transform VL1 with erythromycin selection. The *zip* locus of the resulting strain VL4313, was confirmed by Sanger sequencing.

**Strain VL4315 (D39V, *zip*::$P_3$- BS6-sgRNA3- $P_3$-BS2-*mNeonGreen-opt* (*ery*), $P_{cps}$::$P_3$-BS6-*mScarletI-opt-cps* (*tet*), *cep*:: $P_3$-BS3-sgRNA2 (*spc*), *bgaA*::$P_{lac}$-dCas9-$P_3$-BS3-*mTurquoise2-opt*-$P_3$-BS2-sgRNA6 (*gen*))** was constructed by Golden Gate assembly of three parts. First part is the amplification by PCR of the upstream homologous region to integrate in *S. p. bgaA* locus, the *gen* marker and $P_{lac}$-dCas9 with OVL174 and OVL5899 (with BsaI restriction site) using genomic DNA of strain VL3870 as template. The second part is the amplification by PCR of BssgRNA3-*mTurquoise2-opt*-$P_3$-BS2-sgRNA6 from strain VL3879 with primers OVL5900 and OVL5901 (both with BsaI restriction site). The third part is the amplification by PCR of the downstream homologous region to integrate in *S. p. bgaA* locus from the strain VL3879 with primers OVL5902 (with BsaI restriction site) and OVL173. After purification, the three parts were digested and ligated. The ligation product was used to transform strain VL3875 with gentamycin selection. The resulting strain VL4315 was confirmed by whole genome sequencing (illumina) and also known as the **CAPSUlator**.

**Strain VL4316 (D39V, *zip*:: $P_3$- BS6-sgRNA3- $P_3$-BS2-*mNeonGreen-opt* (*ery*), $P_{cps}$::$P_3$-BS6-*mScarletI-opt-Δcps* (*tet*) (NO *cps*))** was constructed by Golden Gate assembly of two parts. First part is the amplification by PCR of the upstream homologous region to integrate in *S. p. cps* locus, *tetM*, *tetR* and $P_3$-BS6-*mScarletI-opt* from strain VL3872 with primers OVL3689 and OVL5883 (with Esp3I restriction site). The second part is the downstream homologous region to integrate in *S. p. cps* locus from strain VL3872 with primers OVL5156 and OVL5884 (with Esp3I restriction site). After purification, the two parts were digested and ligated together during an assembly reaction with T4 DNA ligase buffer (Vazyme), T4 DNA ligase (Vazyme) and Esp3I (NEB). The assembly mixture was incubated in a thermocycler (PCR Max) for step 1: 1.5 min at 37°C, step 2: 3 min at 16°C, steps 1 and 2 repeated 25x, step 3: 5 min at 37°C and step 4: 10 min at 80°C. Strain VL4313 was directly transformed with the assembly product with tetracycline selection. The *cps* locus of the resulting strain, VL4316, was confirmed by Sanger sequencing.

**Strain VL4317 (D39V, *zip*:: $P_3$- BS6-sgRNA3- $P_3$-BS2-*mNeonGreen-opt* (*ery*), $P_{cps}$::$P_3$-BS6-*mScarletI-opt-Δcps* (*tet*) (NO *cps*), *cep*:: $P_3$-BS3-sgRNA2 (*spc*))** was constructed by amplification of the *cep* locus from gDNA of strain VL4315 with primers OVL873 and OVL874, the PCR product purified was used to transform VL4316 with spectinomycin selection. The *cep* locus of the resulting strain, VL4317, was confirmed by Sanger sequencing.

**Strain VL4318 (D39V, *zip*:: $P_3$- BS6-sgRNA3- $P_3$-BS2-*mNeonGreen-opt* (*ery*), $P_{cps}$::$P_3$-BS6-*mScarletI-opt-cps* (*tet*))** was constructed by amplification of the *cps* locus from gDNA of strain VL4315 with primers

OVL5151 and OVL4901. The purified PCR product was used to transform strain VL4313 with tetracycline selection. The *cps* locus of the resulting strain, VL4318, was confirmed by Sanger sequencing.

**Strain VL4319 (D39V, *zip*:: $P_3$- BS6-sgRNA3- $P_3$-BS2-*mNeonGreen-opt* (*ery*), $P_{cps}$::$P_3$-BS6-*mScarletI-opt-cps* (*tet*), *cep*:: $P_3$-BS3 (*spc*) (NO sgRNA2))** was constructed by Golden Gate assembly of two parts. First part is the amplification by PCR of the upstream homologous region to integrate in the *S. p. cep* locus, the *spc* marker and $P_3$ -BS3 from strain VL3873 with primers OVL873 and OVL5885 (with Esp3I restriction site). Second part is the amplification by PCR of the downstream homologous region to integrate in the *S. p. cep* locus from strain VL3873 with primers OVL5886 (with Esp3I restriction site) and OVL1259. After purification, the two parts were digested and ligated together during an assembly reaction with T4 DNA ligase buffer (Vazyme), T4 DNA ligase (Vazyme) and Esp3I (NEB). The assembly mixture was incubated in a thermocycler (PCR Max) for step 1: 1.5 min at 37°C, step 2: 3 min at 16°C, steps 1 and 2 repeated 25x, step 3: 5 min at 37°C and step 4: 10 min at 80°C. Strain VL4318 was directly transformed with the assembly product with spectinomycin selection. The *cep* locus of the resulting strain, VL4319, was confirmed by Sanger sequencing.

**Strain VL4320 (D39V, *zip*:: $P_3$- BS6-sgRNA3- $P_3$-BS2-*mNeonGreen-opt* (*ery*), *cep*:: $P_3$-BS3-sgRNA2 (*spc*))** was constructed by amplification of the *cep* locus from gDNA of strain VL4315 with primers OVL873 and OVL874. The PCR product was purified and used to transform strain VL4313 with spectinomycin selection. The *cep* locus of the resulting strain, VL4320, was confirmed by Sanger sequencing.

**Strain VL4321 (D39V, *zip*:: $P_3$- BS6-sgRNA3- $P_3$-BS2-*mNeonGreen-opt* (*ery*), $P_{cps}$::$P_3$-BS6-*mScarletI-opt-Δcps* (*tet*) (NO *cps*), *cep*:: $P_3$-BS3-sgRNA2 (*spc*), *bgaA*::$P_{lac}$-dCas9-$P_3$-BS3-*mTurquoise2-opt*-$P_3$-BS2-sgRNA6 (*gen*))** was constructed by PCR amplification of the *bgaA* locus from gDNA of strain VL4315 with primers OVL461 and OVL2871. The purified PCR product was used to transform strain VL4317 with gentamycin selection. The *bgaA* locus of the resulting strain, VL4321, was confirmed by Sanger sequencing and this strain is also referred to as **CAPSUlator-Δ*cps***.

**Strain VL4322 (D39V, *zip*:: $P_3$- BS6-sgRNA3- $P_3$-BS2-*mNeonGreen-opt* (*ery*), $P_{cps}$::$P_3$-BSA6-*mScarletI-opt-cps* (*tet*), *cep*:: $P_3$-BS3 (*spc*) (NO sgRNA2), *bgaA*::$P_{lac}$-dCas9-$P_3$-BS3-*mTurquoise2-opt*-$P_3$-BS2-sgRNA6 (*gen*))** was constructed by PCR of the *bgaA* locus from gDNA of strain VL4315 with primers OVL461 and OVL2871. The PCR product was purified and used to transform strain VL4319 with gentamycin selection. The *bgaA* locus of the resulting strain, VL4322, was confirmed by Sanger sequencing and this strain is also referred to as **CAPSUlator-OFF**.

**Strain VL4323 (D39V, *zip*:: $P_3$- BS6-sgRNA3- $P_3$-BS2-*mNeonGreen-opt* (*ery*), $P_{cps}$::$P_3$-BS1-*mScarletI-opt-cps* (*tet*) (NO BS6 but BS1), *cep*:: $P_3$-BS3-sgRNA2 (*spc*))** was constructed by Golden Gate assembly of two parts. First part is the amplification by PCR of the upstream homologous region to integrate in *S. p. cps* locus, *tetM*, *tetR* and $P_3$ from strain VL3872 with primers OVL5145 and OVL6409 (with SapI restriction site). Second part is the amplification by PCR the downstream homologous region to integrate in *S. p. cps* locus from strain VL3872 with primers OVL6410 (with SapI restriction site) and OVL3692. After purification, the two parts were digested and ligated together during an assembly reaction with T4 DNA ligase buffer (Vazyme), T4 DNA ligase (Vazyme) and SapI (NEB). The assembly mixture was incubated in a thermocycler (PCR Max) for step 1: 1.5 min at 37°C, step 2: 3 min at 16°C, steps 1 and 2 repeated 25x, step 3: 5 min at 37°C and step 4: 10 min at 80°C. Strain VL4320 was directly transformed with the assembly product with tetracycline selection. The *cps* locus of the resulting strain, VL4323, was confirmed by Sanger sequencing.

**Strain VL4324 (D39V, *zip*:: $P_3$- BS6-sgRNA3- $P_3$-BS2-*mNeonGreen-opt* (*ery*), $P_{cps}$::$P_3$-BS1-*mScarletI-opt-cps* (*tet*) (NO BS6 but BS1), *cep*:: $P_3$-BS3-sgRNA2 (*spc*), *bgaA*::$P_{lac}$-dCas9-$P_3$-BS3-*mTurquoise2-opt*-$P_3$-BS2-sgRNA6 (*gen*))** was constructed by amplification by PCR of the *bgaA* locus from gDNA of strain VL4315 with primers OVL461 and OVL2871. The PCR product was purified and used to transform strain VL4323 with gentamycin selection. The *bgaA* locus of the resulting strain, VL4324, was confirmed by Sanger sequencing and this strain is also referred to as **CAPSUlator-ON**.
