## Supplementary material for "Rewiring capsule production by CRISPRi-based genetic oscillators demonstrates a functional role of phenotypic variation in pneumococcal-host interactions": Table S1

**Table S1: Strains and plasmids used in this study**

| Strain | Genotype | Reference |
|---|---|---|
| VL1 | D39V, Serotype 2 strain | [1] |
| VL333 | D39V, Δ*prs1*::$P_{F6}$-*lacI*-*tetR* (*gen*) | [2] |
| VL783 | D39V, *comC::ery* | [3] |
| VL995 | D39V, Δ*prs1*::$P_{F6}$-*lacI* (*gen*), *bgaA*::$P_{lac}$-*dCas9sp* (*tet*), *cil*::$P_3$-*luc* (*kan*), *cep*::$P_3$-*sgRNAluc1* (*spc*) | [4] |
| VL996 | D39V, Δ*prs1*::$P_{F6}$-*lacI* (*gen*), *bgaA*::$P_{lac}$-*dCas9sp* (*tet*), *cil*::$P_3$-*luc* (*kan*), *cep*::pPEPX (*spc*) | [4] |
| VL997 | D39V, Δ*prs1*::$P_{F6}$-*lacI* (*gen*), *bgaA*::$P_{lac}$-*dCas9sp* (*tet*), *cil*::$P_3$-*luc* (*kan*) | [4] |
| VL1998 | DCI23, Δ*prs1*::$P_{F6}$-*lacI* (*gen*), *bgaA*::$P_{lac}$-*dCas9sp* (*tet*) | [4] |
| VL2969 | D39V, Δ*ply*::*camR*, *cil*::*kanR* | Veening lab collection |
| VL3251 | D39V, Δ*prs1*::$P_{F6}$-*lacI* (*gen*), *bgaA*::$P_{lac}$-*dCas9sp* (*tet*), *cil*::$P_3$-*luc* (*kan*), *cep*::$P_3$-BS1-sgRNA*luc* (*spc*) | This study |
| VL3252 | D39V, Δ*prs1*::$P_{F6}$-*lacI* (*gen*), *bgaA*::$P_{lac}$-*dCas9sp* (*tet*), *cil*::$P_3$-*luc* (*kan*), *cep*::$P_3$- BS2-sgRNA*luc* (*spc*) | This study |
| VL3253 | D39V, Δ*prs1*::$P_{F6}$-*lacI* (*gen*), *bgaA*::$P_{lac}$-*dCas9sp* | This study |

| | | |
|---|---|---|
| | (*tet*), *cil*::$P_3$-*luc* (*kan*), *cep*::$P_3$- BS3-sgRNA*luc* (*spc*) | |
| VL3254 | D39V, *Δprs1*::$P_{F6}$-*lacI* (*gen*), *bgaA*::$P_{lac}$-*dCas9sp* (*tet*), *cil*::$P_3$-*luc* (*kan*), *cep*::$P_3$- BS4-sgRNA*luc* (*spc*) | This study |
| VL3255 | D39V, *Δprs1*::$P_{F6}$-*lacI* (*gen*), *bgaA*::$P_{lac}$-*dCas9sp* (*tet*), *cil*::$P_3$-*luc* (*kan*), *cep*::$P_3$- BS5-sgRNA*luc* (*spc*) | This study |
| VL3256 | D39V, *Δprs1*::$P_{F6}$-*lacI* (*gen*), *bgaA*::$P_{lac}$-*dCas9sp* (*tet*), *cil*::$P_3$-*luc* (*kan*), *cep*::$P_3$- BS6-sgRNA*luc* (*spc*) | This study |
| VL3308 | D39V, *Δprs1*::$P_{F6}$-*lacI-tetR* (*gen*), *bgaA*::$P_{lac}$-*dCas9sp* (*tet*) | This study |
| VL3309 | D39V, Δ*prs1*::$P_{F6}$-*lacI* (*gen*), *zip*::$P_{lac}$-*mNeonGreen*-opt-TEV-*mScarletI-opt* (*spc*) | Veening lab collection |
| VL3312 | D39V, Δ*prs1*::$P_{F6}$-*lacI* (*gen*), *zip*::$P_{lac}$-*mNeonGreen-opt*-TEV-*mTurquoise2-opt* (*spc*) | Veening lab collection |
| VL3436 | D39V, *Δprs1*::$P_{F6}$-*lacI-tetR* (*gen*), *bgaA*::$P_{lac}$-*dCas9sp* (*tet*), *cep*::$P_3$-BS3-sgRNA2 (*spc*) | This study |

| | | |
|---|---|---|
| VL3437 | D39V, *Δprs1*::$P_{F6}$-*lacI-tetR* (*gen*), *bgaA*::$P_{lac}$-*dCas9sp* (*tet*), *cep*::$P_3$-BS6-sgRNA3 (*spc*) | This study |
| VL3438 | D39V, *Δprs1*::$P_{F6}$-*lacI-tetR* (*gen*), *bgaA*::$P_{lac}$-*dCas9sp* (*tet*), *cep*::$P_3$-BS2-sgRNA6 (*spc*) | This study |
| VL3439 | D39V, *Δprs1*::$P_{F6}$-*lacI-tetR* (*gen*), *bgaA*::$P_{lac}$-*dCas9sp* (tet), *cep*::$P_3$-BS3-sgRNA2 (*spc*), *zip*::$P_3$-BS2-*mNeonGreen-opt* (*ery*) | This study |
| VL3440 | D39V, *Δprs1*::$P_{F6}$-*lacI-tetR* (*gen*), *bgaA*::$P_{lac}$-*dCas9sp* (*tet*), *cep*::$P_3$-BS3-sgRNA2 (*spc*), *zip*::$P_3$-BS2-*mNeonGreen-opt* (*ery*) | This study |
| VL3441 | D39V, *Δprs1*::$P_{F6}$-*lacI-tetR* (*gen*), *bgaA*::$P_{lac}$-*dCas9sp* (*tet*), cep::$P_3$-BS2-sgRNA6 (*spc*), zip::$P_3$-BS6-*mScarletI-opt* (*ery*) | This study |
| VL3660 | D39V, *Δcps* | Veening lab collection |
| VL3703 | D39V, $P_{cps}$::*tetM-tetR*-$P_{tet}$-*cps* (*tet*) | [2] |
| VL3746 (Strain 1) | D39V, *Δprs1*::$P_{F6}$-*lacI-tetR* (*gen*), *bgaA*::$P_{lac}$-*dCas9sp* (*tet*), *zip*::$P_3$-BS6-sgRNA3-$P_3$-BSA2-*mNeonGreen-opt* (*ery*) | This study |

| | | |
|---|---|---|
| VL3747<br>(Strain 2) | D39V, *Δprs1*::$P_{F6}$-*lacI-tetR* (*gen*), *bgaA*::$P_{lac}$-*dCas9sp* (tet), *cep*::$P_3$-BS6-*mScarletI-opt*-$P_3$-BS3-sgRNA2 (*spc*) | This study |
| VL3748<br>(Strain 3) | D39V, *Δprs1*::$P_{F6}$-*lacI-tetR* (*gen*), *bgaA*::$P_{lac}$-*dCas9sp* (tet), *cil*::$P_3$-BS3-*mTurquoise2-opt*-$P_{tet}$-BS2-sgRNA6 (*kan*) | This study |
| VL3749<br>(Strain 4) | D39V, *Δprs1*::$P_{F6}$-*lacI-tetR* (*gen*), *bgaA*::$P_{lac}$-*dCas9sp* (*tet*), *cil*::$P_3$-BS3-*mTurquoise2-opt*-$P_3$-BS2-sgRNA6 (*kan*) | This study |
| VL3750<br>(Int Strain) | D39V, *Δprs1*::$P_{F6}$-*lacI-tetR* (*gen*), *bgaA*::$P_{lac}$-*dCas9sp* (*tet*), *zip*::$P_3$-BS6-sgRNA3-$P_3$-BS2-*mNeonGreen-opt* (*ery*), *cep*::$P_3$- BS6-*mScarletI-opt*-$P_3$-BS3-sgRNA2 (*spc*) | This study |
| VL3752<br>(Strain 5) | D39V, *Δprs1*::$P_{F6}$-*lacI-tetR* (*gen*), *bgaA*::$P_{lac}$-*dCas9sp* (*tet*), *zip*::$P_3$-BS6-sgRNA3-$P_3$-BS2-*mNeonGreen-opt* (*ery*), *cep*::$P_3$- BS6-*mScarletI-opt*-$P_3$-BS3-sgRNA2 (*spc*), *cil*::$P_3$-BS3-*mTurquoise2-opt*-$P_{tet}$-BS2-sgRNA6 (*kan*) | This study |
| VL3753<br>(Strain 6) | D39V, *Δprs1*::$P_{F6}$-*lacI-tetR* (*gen*), *bgaA*::$P_{lac}$-*dCas9sp* (*tet*), *zip*::$P_3$-BS6-sgRNA3-$P_3$-BS2-*mNeonGreen-opt* (*ery*), | This study |

| | | |
|---|---|---|
| | *cep*::$P_3$- BS6-*mScarletI-opt*-$P_3$-BS3-sgRNA2 (*spc*), *cil*::$P_3$-BS3-*mTurquoise2-opt*-$P_3$-BS2-sgRNA6 (*kan*) | |
| VL3755<br>(Strain5 +AC) | D39V, *Δprs1*:: $P_{F6}$-*lacI-tetR* (*gen*), *bgaA*::Plac-*dCas9*sp (*tet*), *zip*::$P_3$-BS6-sgRNA3-$P_3$-BS2-*mNeonGreen-opt* (*ery*), *cep*::$P_3$-BS6-*mScarletI-opt*-$P_3$-BS3-sgRNA2 (*spc*), *cil*::$P_3$-BS3-*mTurquoise2-opt*-$P_{tet}$-BS2-sgRNA6 (*kan*), *lytA*::*cat*, *comC*::$P_{comC}$-*tmp* | This study |
| VL3757<br>(Strain6 +AC) | D39V, *Δprs1*::$P_{F6}$-*lacI-tetR* (*gen*), *bgaA*::$P_{lac}$-*dCas9sp* (*tet*), *zip*::$P_3$-BS6-sgRNA3-$P_3$-BS2-*mNeonGreen-opt* (*ery*), *cep*::$P_3$- BS6-*mScarletI-opt*-$P_3$-BS3-sgRNA2 (*spc*), *cil*::$P_3$-BS3-*mTurquoise2-opt*-$P_3$-BS2-sgRNA6 (*kan*), *lytA::cat*, *comC*::$P_{comC}$-*tmp* | This study |
| VL3869<br>(VL3755+del-cps) | D39V, *Δprs1*:: $P_{F6}$-*lacI-tetR* (*gen*), *bgaA*::Plac-*dCas9*sp (*tet*), *zip*::$P_3$-BS6-sgRNA3-$P_3$-BS2-*mNeonGreen-opt* (*ery*), *cep*::$P_3$-BS6-*mScarletI-opt*-$P_3$-BS3-sgRNA2 (*spc*), *cil*::$P_3$-BS3-*mTurquoise2-opt*-$P_{tet}$-BS2-sgRNA6 (*kan*), *lytA::cat*, *comC*::$P_{comC}$-*tmp, Δcps* | This study |

| | | |
|---|---|---|
| VL3870 | D39V, *bgaA::*$P_{lac}$-*dcas9sp* (*gen*) | This study |
| VL3872 | D39V, $P_{cps}$::*tetM*-*tetR*-$P_3$-BS6-*mScarletI-opt-cps* (*tet*) | This study |
| VL3873 | D39V, $P_{cps}$::*tetM*-*tetR*-$P_3$-BssgRNA6-*mScarletI-opt-cps* (*tet*), *cep*::$P_3$-BS3-sgRNA2 (*spc*) | This study |
| VL3875 | D39V, $P_{cps}$::*tetM*-*tetR*-$P_3$-BS6-*mScarletI-opt-cps* (*tet*), *cep*::$P_3$-BS3-sgRNA2 (*spc*), *zip*::$P_3$-BS6-sgRNA3-$P_3$-BS2-*mNeonGreen-opt* (*ery*) | This study |
| VL3876 | D39V, $P_{cps}$::*tetM*-*tetR*-$P_3$-BS6-*mScarletI-opt-cps* (*tet*), *cep*::$P_3$-BS3-sgRNA2 (*spc*), *zip*::$P_3$-BS6-sgRNA3-$P_3$-BS2-*mNeonGreen-opt* (*ery*), *cil*::$P_3$-BS3-*mTurquoise2-opt*-$P_{tet}$-BS2-sgRNA6 (*kan*) | This study |
| VL3877 | D39V, $P_{cps}$::*tetM*-*tetR*-$P_3$-BS6-*mScarletI-opt-cps* (*tet*), *cep*::$P_3$-BS3-sgRNA2 (*spc*), *zip*::$P_3$-BS6-sgRNA3-$P_3$-BS2-*mNeonGreen-opt* (*ery*), *cil*::$P_3$-BS3-*mTurquoise2-opt*-$P_3$-BS2-sgRNA6 (*kan*) | This study |
| VL3878 | D39V, $P_{cps}$::*tetM*-*tetR*-$P_3$-BS6-*mScarletI-opt-cps* (*tet*), *cep*::$P_3$-BS3-sgRNA2 (*spc*), *zip*::$P_3$- | This study |

| | | |
|---|---|---|
| | BS6-sgRNA3-$P_3$-BS2-*mNeonGreen-opt* (*ery*), *cil*::$P_3$-BS3-*mTurquoise2-opt*-$P_{tet}$-BS2-sgRNA6 (*kan*), *bgaA*::$P_{lac}$-*dcas9sp* (*gen*)<br><br>Inducible CAPSUlator 1.0 | |
| VL3879 | D39V, $P_{cps}$::*tetM-tetR*-$P_3$-BS6-*mScarletI-opt-cps* (*tet*), *cep*::$P_3$-BS3-sgRNA2 (*spc*), *zip*::$P_3$-BS6-sgRNA3-$P_3$-BS2-*mNeonGreen-opt* (*ery*), *cil*::$P_3$-BS3-*mTurquoise2-opt*-$P_3$-BS2-sgRNA6 (*kan*), *bgaA*::$P_{lac}$-*dcas9sp* (*gen*)<br><br>Constitutive CAPSUlator 1.0 | This study |
| VL3882 | D39V, *Δprs1::$P_{F6}$-lacI (gen), zip::tmp-$P_{lac}$-MCS (tmp)* | Veening lab collection |
| **VL4313** | D39V, *zip*:: $P_3$- BS6-sgRNA3- $P_3$-BS2-*mNeonGreen-opt* (*ery*) | This study |
| VL4315 | D39V, *zip*:: $P_3$- BS6-sgRNA3- $P_3$-BS2-*mNeonGreen-opt* (*ery*), $P_{cps}$::*tetM-tetR*-$P_3$-BS6-*mScarletI-opt-cps* (*tet*), *cep*:: $P_3$-BS3-sgRNA2 (*spc*) , *bgaA*::$P_{lac}$-dCas9-$P_3$-BS3-*mTurquoise2-opt*-$P_3$-BS2-sgRNA6 (*gen*)<br><br>CAPSUlator 2.0 | This study |
| VL4316 | D39V, *zip*:: $P_3$- BS6-sgRNA3- $P_3$-BS2-*mNeonGreen-opt* (*ery*), | This study |

| | | |
|---|---|---|
| | $P_{cps}$::*tetM-tetR*-$P_3$-BS6-*mScarletI-opt*-Δ*cps* (*tet*) (NO *cps*) | |
| VL4317 | D39V, *zip*:: $P_3$- BS6-sgRNA3- $P_3$-BS2-*mNeonGreen-opt* (*ery*), $P_{cps}$::*tetM-tetR*-$P_3$-BS6-*mScarletI-opt*-Δ*cps* (*tet*) (NO *cps*), *cep*:: $P_3$-BS3-sgRNA2 (*spc*) | This study |
| VL4318 | D39V, *zip*:: $P_3$- BS6-sgRNA3- $P_3$-BS2-*mNeonGreen-opt* (*ery*), $P_{cps}$::*tetM-tetR*-$P_3$-BS6-*mScarletI-opt-cps* (*tet*) | This study |
| VL4319 | D39V, *zip*:: $P_3$- BS6-sgRNA3- $P_3$-BS2-*mNeonGreen-opt* (*ery*), $P_{cps}$::*tetM-tetR*-$P_3$-BS6-*mScarletI-opt-cps* (*tet*), *cep*:: $P_3$-BS3 (*spc*) (NO sgRNA2) | This study |
| VL4320 | D39V, *zip*:: $P_3$- BS6-sgRNA3- $P_3$-BS2-*mNeonGreen-opt* (*ery*), *cep*:: $P_3$-BS3-sgRNA2 (*spc*) | This study |
| VL4321 | D39V, *zip*:: $P_3$- BS6-sgRNA3- $P_3$-BS2-*mNeonGreen-opt* (*ery*), $P_{cps}$::*tetM-tetR*-$P_3$-BS6-*mScarletI-opt*-Δ*cps* (*tet*) (NO *cps*), *cep*:: $P_3$-BS3-sgRNA2 (*spc*), *bgaA*::$P_{lac}$-dCas9-$P_3$-BS3-*mTurquoise2-opt*-$P_3$-BS2-sgRNA6 (*gen*)<br>CAPSU2-Ctl-Delta-cps | This study |

| | | |
|---|---|---|
| VL4322 | D39V, *zip*:: $P_3$- BS6-sgRNA3- $P_3$-BS2-*mNeonGreen-opt* (*ery*), $P_{cps}$::*tetM*-*tetR*-$P_3$-BS6-*mScarletI-opt-cps* (*tet*), *cep*:: $P_3$-BS3 (*spc*) (NO sgRNA2), *bgaA*::$P_{lac}$-dCas9-$P_3$-BS3-*mTurquoise2-opt*-$P_3$-BS2-sgRNA6 (*gen*)<br>CAPSU2-Ctl-Always-OFF | This study |
| VL4323 | D39V, *zip*:: $P_3$- BS6-sgRNA3- $P_3$-BS2-*mNeonGreen-opt* (*ery*), $P_{cps}$::*tetM*-*tetR*-$P_3$-BS1-*mScarletI-opt-cps* (*tet*) (NO BS6 but BS1), *cep*:: $P_3$-BS3-sgRNA2 (*spc*) | This study |
| VL4324 | D39V, *zip*:: $P_3$- BS6-sgRNA3-$P_3$-BS2-*mNeonGreen-opt* (*ery*), $P_{cps}$::*tetM*-*tetR*-$P_3$-BS1-*mScarletI-opt-cps* (*tet*) (NO BS6 but BS1), *cep*::$P_3$-BS3-sgRNA2 (*spc*), *bgaA*::$P_{lac}$-dCas9-$P_3$-BS3-*mTurquoise2-opt*-$P_3$-BS2-sgRNA6 (*gen*)<br>CAPSU2-Ctl-Always-ON | This study |
| **Plasmid** | **Genotype** | **Reference** |
| pVL1305 | pPEPX-$P_3$-sgRNA*luc, spc* | Veening lab collection |
| pASR102 | pPEPZ-$P_{lac}$, *gen* | Keller et al |
| pASR103 | pPEPZ-$P_{lac}$, *ery*, *bla* | Keller et al |

| | | |
|---|---|---|
| pASR110 | pPEPZ-$P_{lac}$-*mNeonGreen-opt*, *spc* | Veening lab collection |
